# Mapping current and future thermal limits to suitability for malaria transmission by the invasive mosquito *Anopheles stephens*i

**DOI:** 10.1101/2022.12.15.520598

**Authors:** Sadie J. Ryan, Catherine A. Lippi, Oswaldo C. Villena, Aspen Singh, Courtney C. Murdock, Leah R. Johnson

## Abstract

**Background:** *Anopheles stephensi* is a malaria-transmitting mosquito that has recently expanded from its primary range in Asia and the Middle East, to locations in Africa. This species is a competent vector of both *P. falciparum* (PF) and *P. vivax* (PV) malaria. Perhaps most alarming, the characteristics of *An. stephensi*, such as container breeding and anthropophily, make it particularly adept at exploiting built environments in areas with no prior history of malaria risk.

**Methods:** In this paper we created global maps of thermal transmission suitability and people at risk (PAR) for malaria transmission by *An. stephensi*, under current and future climate. Temperature-dependent transmission suitability thresholds derived from recently published species-specific thermal curves were used to threshold gridded, monthly mean temperatures under current and future climatic conditions. These temperature driven transmission models were coupled with gridded population data for 2020 and 2050, under climate-matched scenarios for future outcomes, to compare with baseline predictions for 2020 populations.

**Results:** Using the Global Burden of Disease regions approach, we found that heterogenous regional increases and decreases in risk did not mask the overall pattern of massive increases of PAR for malaria transmission suitability with *An. stephensi* presence. General patterns of poleward expansion for thermal suitability were seen for both PF and PV transmission potential.

**Conclusions:** Understanding the potential suitability for *An. stephensi* transmission in a changing climate provides a key tool for planning, given an ongoing invasion and expansion of the vector. Anticipating the potential impact of onward expansion to transmission suitable areas, and the size of population at risk under future climate scenarios, and *where* they occur, can serve as a large-scale call for attention, planning, and monitoring.

## Background

Malaria remains a critical global health challenge, with 241 million cases reported by the World Health Organization (WHO) in 2020 alone [1]. Although the current distribution of malaria is largely pantropical, the overwhelming majority of cases and deaths occur in non-arid regions of Africa, where WHO estimated approximately 600,000 deaths in 2020 [1–3]. Malarial transmission throughout sub-Saharan Africa is historically attributable to a few key mosquito vectors, most notably those in the *Anopheles gambiae* species complex [1,4]. However, in 2019 the WHO issued a notice to alert public health authorities to the recent expansion of invasive *Anopheles stephensi* into the Horn of Africa, identifying this new vector species as a major potential threat to malaria control in the region [5–7].

The expansion of *An. stephensi* represents a new critical threat not only to communities in Africa, but also to global public health. A competent vector of both *Plasmodium falciparum* (PF) and *P. vivax* (PV) malaria, *An. stephensi* has been implicated in malaria transmission throughout much of its native range in Asia and the Middle East, including India, Iran, and Pakistan [8–12]. In contrast with other Anopheline species, *An. stephensi* is capable of exploiting containers of standing water for ovipositional habitat, similar to container-breeding mosquitoes in the genus *Aedes*, including *Ae. aegypti* and *Ae. albopictus* [13]. This notable difference in life history has enabled the incursion of *An. stephensi* into built environments, fueling urban outbreaks of malaria and facilitating invasions into new geographic areas. In the past decade, *An. stephensi* has successfully expanded its range into the African continent, with established populations in Djibouti, Ethiopia, and Sudan [14]. Alarmingly, the arrival of this new vector has precipitated epidemics in populations centrally located in urban areas, where rates of malaria have historically been significantly lower compared to rural and peri-urban areas [15]. This shift in underlying risk was exemplified by a notable malaria outbreak in Djibouti City in 2012, where such outbreaks have since become increasingly severe, and are now an annual occurrence [16,17]. Other physiological adaptations of *An. stephensi*, such as acquired insecticide resistance and greater range of thermal tolerance compared to *An. gambiae*, raise further concerns regarding the continued success of this mosquito as an invasive species, and its ability to potentially undermine existing vector control strategies [18–20]. Perhaps unsurprisingly, *An. stephensi* has been identified as a major risk to malaria elimination targets, with global public health organizations calling for increased entomological surveillance and vector control in areas at imminent risk of invasion [5].

Mapping geographic estimates of transmission suitability can provide essential tools for assessing the current and future risk of *An. stephensi* invasions, and subsequent malaria transmission. In previous papers, we mapped malaria suitability for Africa, using a model comprised of an array of *Anopheles spp*. input parameters, primarily for *P. falciparum* malaria transmitted by *An. gambiae* [21]. In a recently updated model, *An. gambiae* and *An. stephensi,*and the two main malarial parasites they transmit (PF and PV), were separately modeled to produce thermal suitability curves for transmission [20,21]. While the potential geographic dispersal of *An. gambiae* is functionally limited by arid conditions, *An. stephensi* is a container breeder that is resilient to habitat extremes [5]. Therefore, *An. stephensi* is able to thrive in close association with people, and thus potentially able to establish itself everywhere that temperature is not limiting. Thus, understanding the potential areas for suitability for transmission by this invasive mosquito now, and in the future, is important for capacity building and planning control efforts.

In this study, we map the global suitability of malaria transmission by *An. stephensi*, using modeled thermal limits under current and future climate scenarios. Unlike previous studies to map the distribution of malaria, we are not limiting the projected distributions to non-arid regions, given the life history of *An. stephensi*, and instead make similar assumptions to those for mapping *Aedes* spp. transmitted diseases. Additionally, *An. stephensi* thermal suitability maps were combined with projected human population density estimates, enabling us to assess not only the areas that are vulnerable to malaria transmission through *An. stephensi* expansions, but also the magnitude of threat in terms of people at risk (PAR).

## Methods

### Thermal suitability model

In a previous study, we used mechanistic modeling to establish thermal suitability curves for transmission of PF and PV by *An. stephensi*[20]. Briefly, thermal response data for vector and parasite pairings were synthesized from published data. These data were used to parameterize a formulation for *R_0_*, the basic reproductive number, that explicitly incorporated temperature-dependent traits for both mosquito vectors and malarial parasites. The temperature-dependent components of the *R_0_* formulation were used to define a suitability metric *S(T)*, defined as:

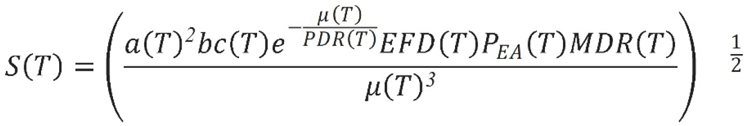

Where *a* is mosquito biting rate; *bc* is vector competence; μ is the mosquito mortality rate; *PDR* is the parasite development rate; EFD is mosquito fecundity expressed as the number of eggs per female per day; *P_EA_* is the proportion of eggs surviving to adulthood; and *MDR* is the mosquito development rate.

A Bayesian approach was used to fit unimodal thermal response curves of traits for each mosquito or parasite species. Samples from the resulting joint posterior distribution of the suitability metric were used to calculate overall thermal response, in addition to critical temperature thresholds for pathogen transmission by species [20].

In this study we used the thermal boundaries where the malaria transmission suitability metric for *An. stephensi* was greater than zero (S(T) >0), with a posterior probability greater than 0.975 [20]. The resulting thermal limits for malaria transmitted by *An. stephensi* are temperatures of 16.0-36.5 °C for PF and 16.6 - 31.7 °C for PV *[20].*

### Climate data

In this paper, baseline and future scenarios for *An. stephensi* transmitted PF and PV suitability are described. Outputs are presented for a baseline climate scenario, and future potential climate driven outputs for four General Circulation Models (GCMs), following the methodology used in Ryan et al. 2019 and Ryan et al. 2021 [22,23] to describe climate impacts on the global distribution of *Aedes* spp. transmitted diseases.

Baseline and future scenario climate model output data were acquired from the research program on Climate Change, Agriculture, and Food Security (CCAFS) web portal (http://ccafs-climate.org/data_spatial_downscaling/), part of the Consultative Group for International Agricultural Research (CGIAR). The baseline climate model from which these are projected is the WorldClim v1.4 baseline [24], and thus it serves as our baseline for our models. The CCAFS future model outputs were created using the delta downscaling method, from the IPCC AR5. The GCMs used in this study are the Beijing Climate Center Climate System Model (BCC-CSM1.1); the Hadley GCM (HadGEM2-AO and HadGEM2-ES); and the National Center for Atmospheric Research’s Community Climate System Model (CCSM4). The datasets were obtained at a resolution of 5-arc minutes, matching the spatial resolution of baseline data.

For visualizations, we used one of the General Circulation Models (GCMs): the Hadley Centre Global Environment Model version 2, Earth-System configuration (HADGEM2-ES) under two scenarios for greenhouse gas emissions, or representative concentration pathways (RCPs): RCP 4.5 and RCP 8.5. Mechanistic transmission models were projected onto climate data in R (v. 4.1.2) with the package ‘raster’[25]. Monthly mean temperatures were thresholded according to the thermal suitability limits for each malaria species, and the number of suitable months of transmission summed (0-12) in a pixel-wise analysis for the globe.

### Population data

In order to establish a population baseline, the 2020 Gridded Population of the World (GPW4, ver 4.11) was used [26]. The decision about how to best match climate baselines with population is complex, as baselines represent climate normal periods around the start of the 21st century, rather than a ‘current’ climate baseline. However, we chose the nearest decade to current conditions for population baseline. For the future population, we chose to use 2050 projections for two Shared Socioeconomic Pathways (SSPs) [27,28], best matched to the RCPs we chose. As the combinations of RCPs and SSPs are not all realistic, CMIP5 RCP 4.5 and RCP 8.5 for SSP2 and SSP5, respectively were modeled. We aggregated all geographic layers in our analyses to a 0.25° grid cell for consistency.

Monthly suitability maps were produced for baseline and future scenarios, using one GCM (HADGEM-ES), for illustration (Figure 1). Maps were created using ArcGIS ver 10.1[29].

**Figure 1.**
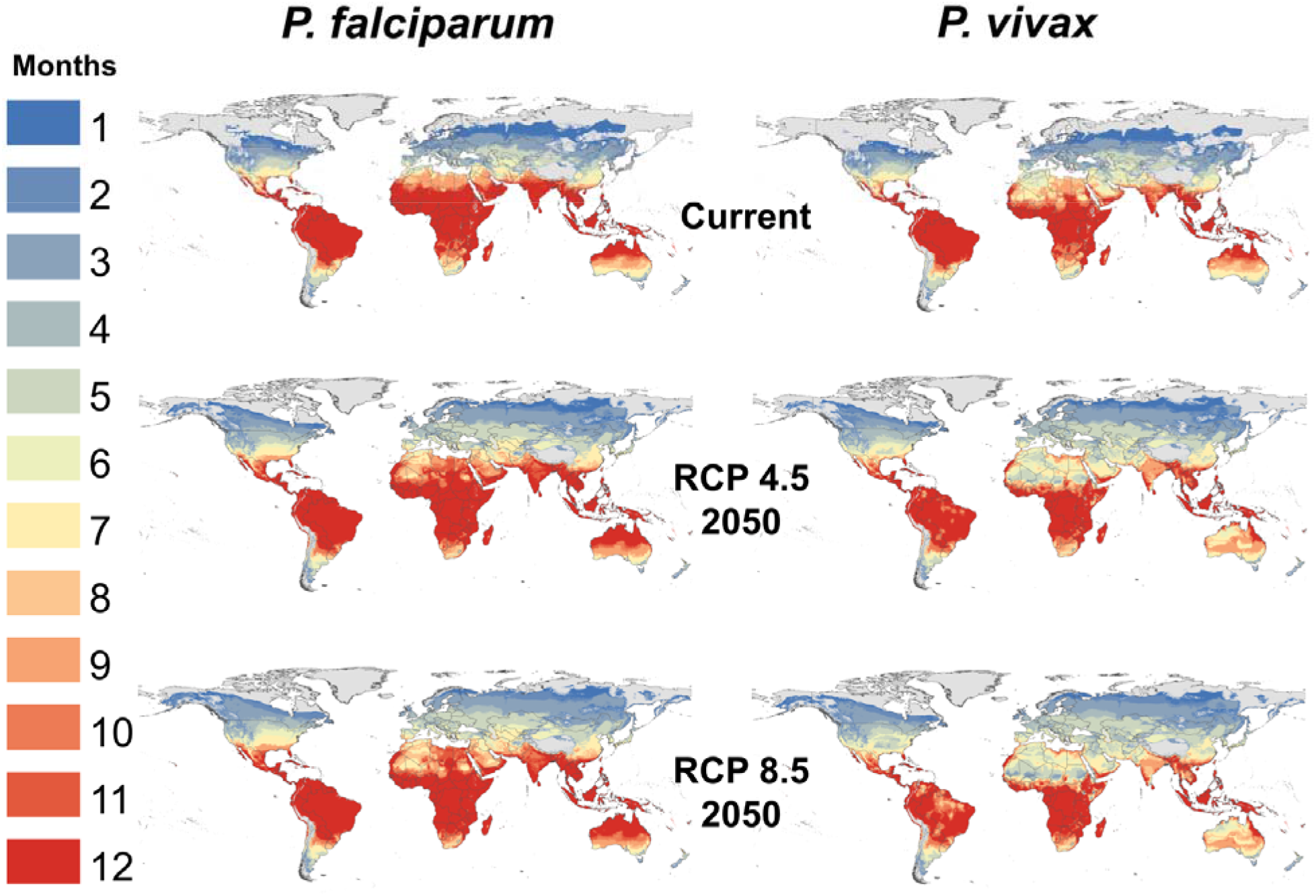
Thermal suitability for transmission of *P. falciparum* and *P. vivax* malaria by *An. stephensi.*Transmission suitability is shown under current climate conditions, and for the year 2050 at RCP 4.5 and RCP 8.5. The number of months of suitable temperatures are given as shaded areas, where the posterior probability of S(T)>0 is 0.975.

To describe the impact to populations in current and future landscapes, we used the Global Burden of Disease (GBD) regions [30] to summarize the population at risk (PAR). Tables S1–4 summarize the top 10 regions, and global gain overall, in terms of increased (difference between current and projected) PAR for both year-round (12 month, endemic), and for ‘any’ (one or more months), for each of transmission suitability of PF and PV by *An. stephensi*, under the two future RCP × SSP scenarios, averaged across the 4 GCMs (also summarized at a global level in Figure 2a,b).

**Figure 2a:**
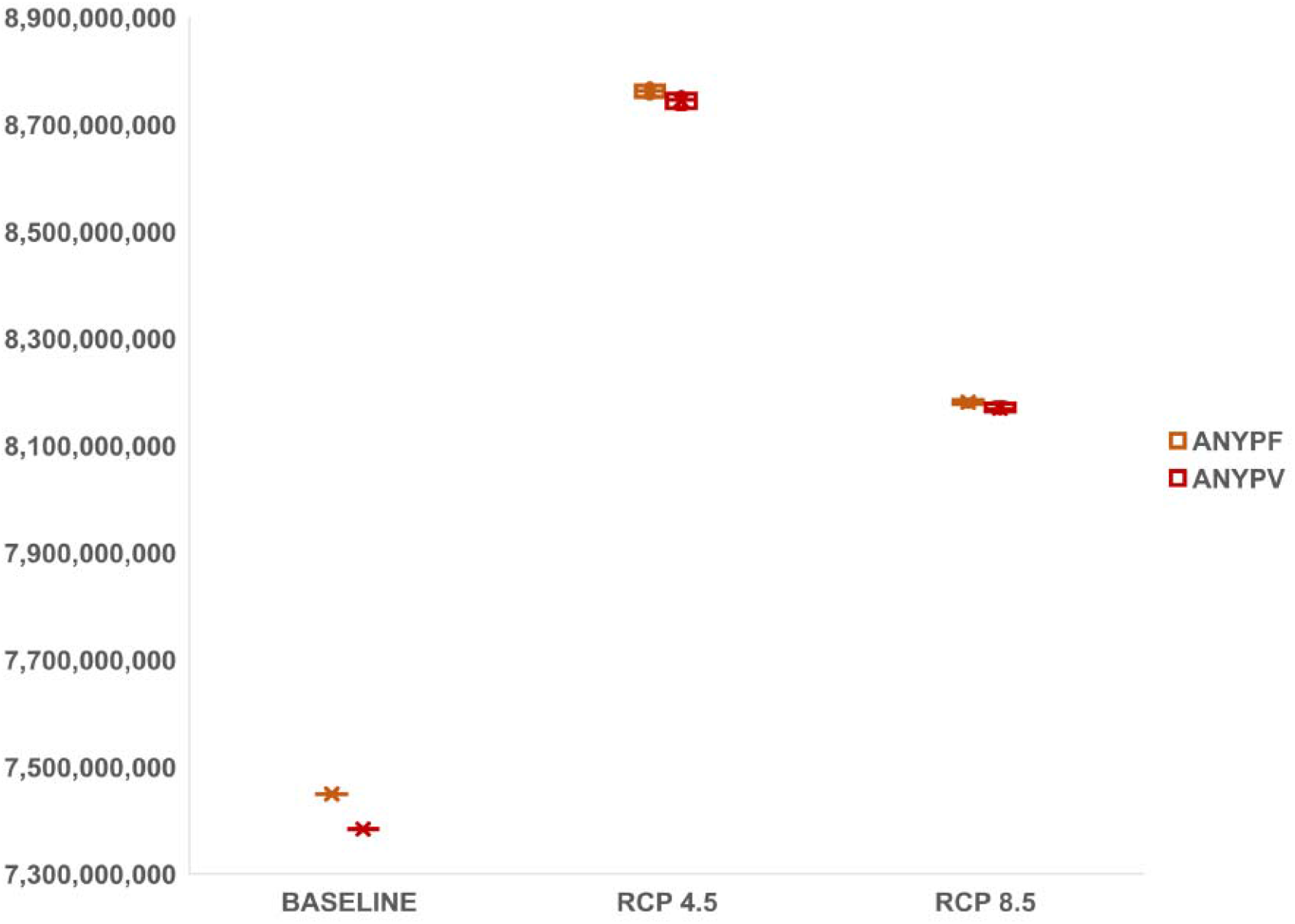
Future projected global (net) population at risk (PAR) for any (one or more months) transmission suitability for *P. falciparum* (orange) and *P. vivax* (red) by *An. stephensi*, under RCP 4.5 (SSP2) and RCP 8.5 (SSP5) scenarios, across four GCMs (shown as box and whiskers here, mean markers included (x)).

**Figure 2b:**
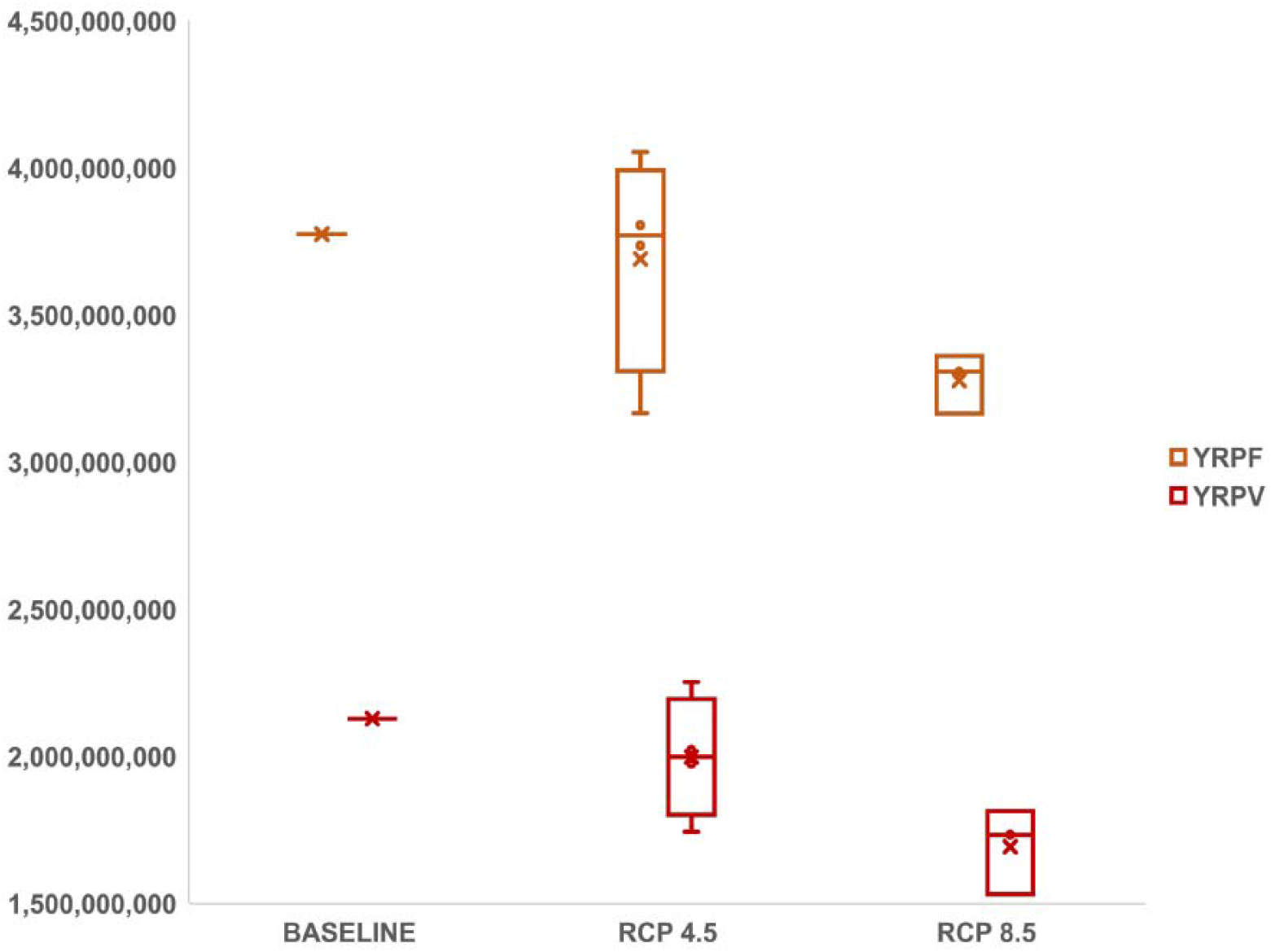
Future projected global (net) population at risk (PAR) for year round (12 months, endemic) transmission suitability for *P. falciparum* (orange) and *P. vivax* (red) by *An. stephensi*, under RCP 4.5 (SSP2) and RCP 8.5 (SSP5) scenarios, across four GCMs (shown as box and whiskers here, mean markers included (x)).

## Results

### Baseline suitability and duration of transmission season

Maps of months of *An. stephensi* malaria transmission suitability for PF and PV are shown in Figure 1. Much of Africa is projected to be suitable at baseline for nearly year-round transmission for both malaria parasites. Beyond Africa, the predicted baseline thermal transmission suitability for both PF and PV extends throughout the global tropics, throughout the known existing range of the Middle East, extending throughout Asia, Central America, South America, and marginally in North America. The predicted potential range for year-round transmission suitability of PF extends further North and South than PV, with notable extension of the transmission season in northern Africa, the Middle East, India, and central Australia. The narrower thermal suitability bounds for PV constrains the baseline potential extent, compared to that for PF. Seasonal transmission suitability at baseline climate conditions is projected to be globally widespread for both PF and PV, extending well into temperate regions in North America, Europe, and Asia.

### Predicted future suitability

Mapped transmission suitability in 2050, for RCP 4.5 and RCP 8.5 scenarios is shown in Figure 1. We see an expansion of the transmission suitability season for PF in both RCPs. Notably, the potential for any transmission (i.e., one or more months) is predicted to expand at northern latitudes, where areas with no current malaria suitability will have the potential for transmission, at least for one month every year. This includes portions of Alaska in the US, northern Canada, Scandinavia, and Russia. The length of the PF transmission season is expected to increase in temperate regions of North America and Europe, southern Africa, and in central Australia. Yet, changing climate conditions will shorten the length of the PF transmission season in some areas, most notably in northern Africa, the Middle East, and northern India. The transmission suitability for PV is also predicted to extend further North in the future, encroaching on areas that do not currently experience malaria transmission. The length of the PV transmission season is expected to increase in southern Africa and in parts of North America, including Mexico and along the Gulf Coast in the US. There are also marked decreases in the length of the PV transmission season, most notably throughout northern Africa, the Middle East, India, Asia, northern Australia, and South America.

### Population at risk

The projections of thermal suitability for transmission of PF and PV by AS for the two future scenarios of RCP 4.5 and RCP 8.5, in combination with matched population projections SSP2 and SSP5 respectively, revealed that in many regions of the world, increases in people at risk of transmission suitability will occur (Tables S1–S4). As the prediction for SSP2 population for the globe is larger than SSP5 in 2050 (9.17 billion vs 8.56 billion [31]), our results reflect the combination of potential geographic shifts of suitability and the underlying population changes. Perhaps counterintuitively, our RCP 4.5 scenario predicts a larger net increase than our RCP 8.5, for PAR in both ‘any’ (one or more) transmission, and year-round (endemic) transmission scenarios (Figure 2a,b). The baseline and net global future population at risk (PAR) for transmission suitability is shown across the 4 GCMs used in this study in Figure 2, as boxplots, comparing PF and PV suitability.

At 2020 population baseline, 7.45 billion people are predicted to be at risk for transmission suitability for one or more months for PF in *An. stephensi*, and 7.38 billion for PV in AS. Under RCP 4.5, the net PAR for PF suitability increases to a range of 8.75-8.77 billion, and for PV 8.73-8.76 billion, across the 4 GCMs; and under RCP 8.5, the estimated PAR for PF suitability is 8.18-8.19 billion, and for PV 8.16-8.18 billion (Figure 2a).

At baseline, the year round PAR for PV is 2.13 billion people, while it is 3.77 billion for PF, emphasizing the difference in risk imposed by the broader temperature range of suitability for PF. Under RCP 4.5, the net PAR for PF suitability increases to a range of 3.73-4.05 billion, and 1.98-2.25 billion for PV. Under RCP 8.5 conditions, net par for PF suitability increases to 3.16-3.36 billion, and 1.53-1.81 billion for PV.

The top 10 largest regional increases in PAR for each of PF, PV, and for year-round and ‘any’ transmission are given in Tables S1–4, including the global gains in increases (in contrast to net changes). For PF, for endemic (year-round) transmission PAR increases, East and West Sub-Saharan Africa regions are the top two affected, under both the RCP 4.5 and RCP 8.5 scenario (Table S1); for PV, while East Sub-Saharan Africa is also the top affected region, the second place is Central African Region, indicating a shifted geographic impact with the narrower thermal bounds (Table S2). For ‘any’ (one or more months) transmission suitability, South Asia region is the top affected for both PF and PV, ahead of East Sub-Saharan Africa region, suggesting a shift of seasonal, sub-endemic risk into high density population areas in both of the future scenarios explored here.

## Discussion

Assessing the future risk of *An. stephensi* expansion against the backdrop of changing climate is imperative for public health planning and risk mitigation. Its propensity to spread and establish outside of its current range is already underway, drawing the attention of the global health community [5,7,17,32]. As a malaria transmitting Anopheline capable of exploiting a similar niche to the urban adapted *Aedes* spp. mosquitoes, anticipating where the bounds of thermal limits to persistence and transmission exist is a step towards understanding where it can invade and establish, both now and in the future.

Using a recently published thermal suitability model for transmission of both PF and PV malaria by *An. stephensi*[20], mapped months of suitability demonstrated that a large part of the world is already suitable for one or more months of the year, putting an estimated 7.38-7.45 billion people at risk of that potential. While the actual arrival, establishment, and onwards transmission of malaria may be less risky for areas with a low number of months of suitability, this approach indicates that a baseline of 2.13-3.77 billion people are currently living in places with endemic risk - not simply in the known existing range for transmission by *An. stephensi*. The potential for future shifts in the range of suitable areas was explored, as a function of a changing climate and shifting population projections, reflective of those potential climate scenarios. The mapped number of months of transmission suitability showed a poleward expansion of areas becoming suitable, and a shift from some lower latitude locations to becoming hotter than suitable for transmission during parts of the year, shortening the season. While this shift away from suitability results in predicted declines in risk to populations, as transmission suitability shifts out of hotter regions, other health crises are exacerbated at overly high temperatures [33–36], and this is thus not cause for less alarm, nor is it mitigation for malaria.

This exploration of potential future climate impacts on a vector currently expanding its range is based on current vector-pathogen biology and thermal limits to the life-history of *An. stephensi* and the malaria parasites. The conditions under which parameters in the underlying thermal suitability model were established were idealized laboratory conditions, and are not yet established for *An. stephensi* undergoing the climate changes we model here. The environment experienced in a changed climate in 2050 may induce different interactions between vectors and their pathogens, but we anticipate that the plastic responses of the vector (e.g. urban adaptation, behavioral avoidance of environmental extremes) and the pathogen (e.g. rapid evolution under novel environmental pressures, or fluctuating temperatures), and how that will impact the vector microbiome [37], potentially altering vector competence, will lead to broader potential temperature limits to suitability, making the estimates presented here conservative.

A knock-on effect of the potential expansion of a novel, urban-adapted, malaria vector into, for example, the Americas, is that adding a competent malaria vector to areas with existing competent malaria vectors expands the competent vector community. This compounds the risk in a changing world for facilitating spillover from a novel invader experiencing perhaps only a shortened suitability season to established Anopheline species (e.g. *An. quadrimaculatus* in parts of N. America).

The ongoing expansion of *An. stephensi* is troubling, given its implication in the shift from primarily rural to urban malaria transmission. Although urban malaria transmission represents a new threat for many existing vector control programs to manage, there may be opportunities for the formation of successful mitigation efforts, given that agencies are aware of potential expansion. With enough lead time, mosquito control agencies may be able to successfully leverage knowledge, experience, and tools for controlling other urban container-breeding mosquitoes to suppress the proliferation of invasive *An. stephensi* [6]. For example, dengue fever surveillance and control programs that target *Ae. aegypti* may have the capacity to expand efforts to include *An. stephensi* without a major investment in novel resources. Though there are still challenges in program adaptation, such as the need to address insecticide resistance, it is likely that effective control measures will not have to be designed from the ground up.

## Conclusion

Mapping thermal suitability for malaria transmission for the invasive urban-adapted *An. stephensi* for baseline and future climate and population projection scenarios shows that much of the world is suited to continued range expansion now and into the future. While this work demonstrates that around a third of the world’s population lives in areas of potential risk, understanding where range expansion is plausible, and how that may shift in the future, provides broad scale tools for motivating surveillance and opportunities for preemptive interventions. Of key importance, the similarity between *An. stephensi* and *Aedes spp*, and their management as urban container breeders may provide an opportunity to leverage existing vector management and control for *An. stephensi*.

## List of Abbreviations

CGIAR: Consultative Group for International Agricultural Research
CMIP: Climate Model Intercomparison Project
GCM: General Circulation Model
GPW: Gridded Population of the World
RCP: Representative Concentration Pathway
SSP: Shared Socioeconomic Pathway
WHO: World Health Organization

## Declarations

### Ethics approval and consent to participate

Not applicable

### Consent for publication

Not applicable

### Availability of data and materials

All model output rasters are available through Harvard Dataverse (URL - to be completed per journal requirements), all climate layers are freely available online and described within the paper. R Code for creating model outputs available (to be completed per journal requirements)

### Authors’ contributions

SJR, CAL and AS conducted analyses; SJR and CAL drafted the paper, figures and tables; all authors contributed to the final version.

## Acknowledgements

The authors would like to acknowledge the thoughtful insights of our anonymous reviewers.

---

**Table S1.**
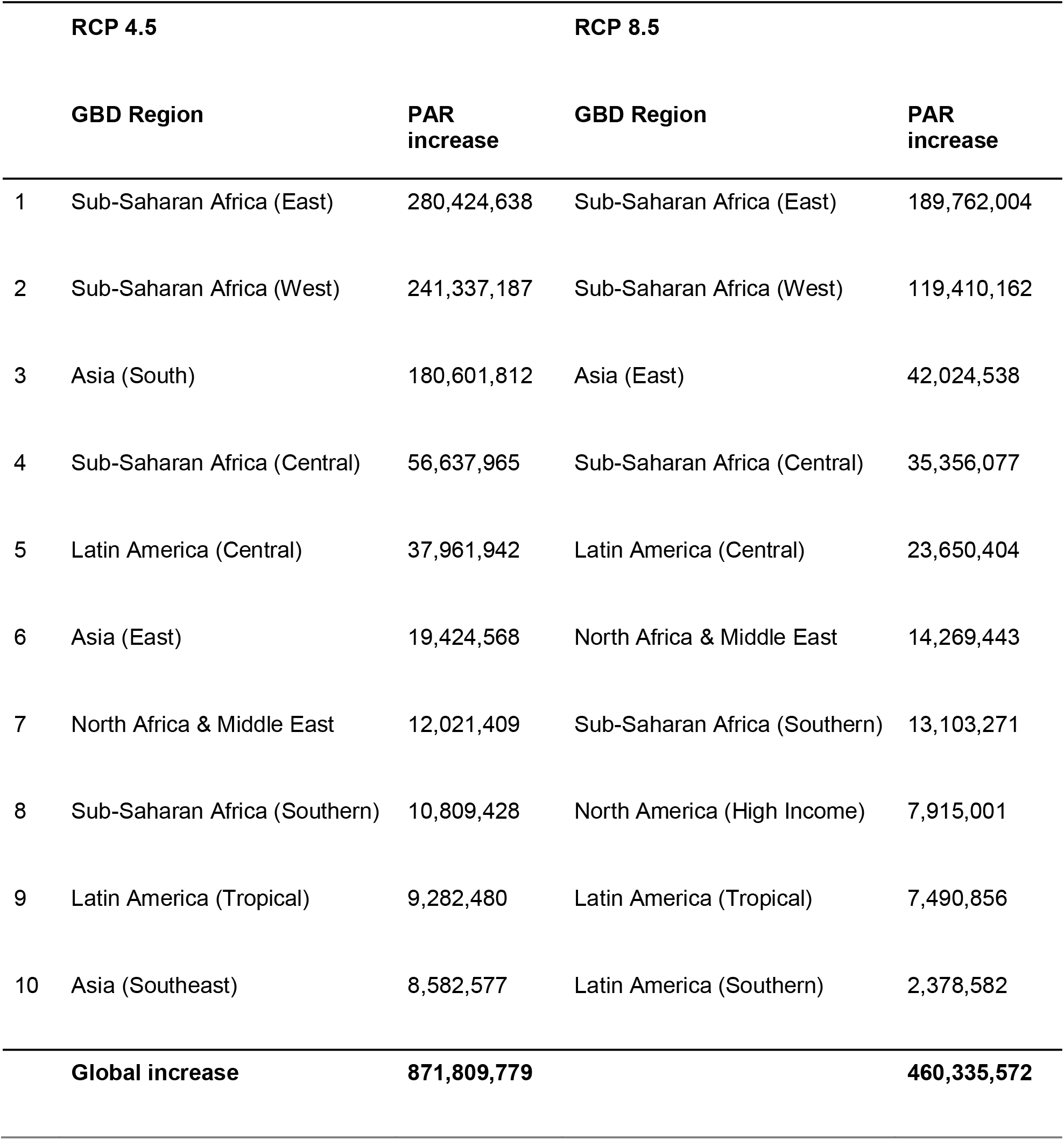
Top 10 Global Burden of Disease defined regional increase in people at risk (PAR) for year-round transmission suitability of *P. falciparum* by *An. stephensi* in 2050, under RCP 4.5 (SSP2 population projection) and RCP 8.5 (SSP5 population projection) future climate scenarios, averaged across four general circulation models (GCMs) as described in the main methods. Global increase is the sum of all gains in PAR increases across all GBD regions.

**Table S2.**
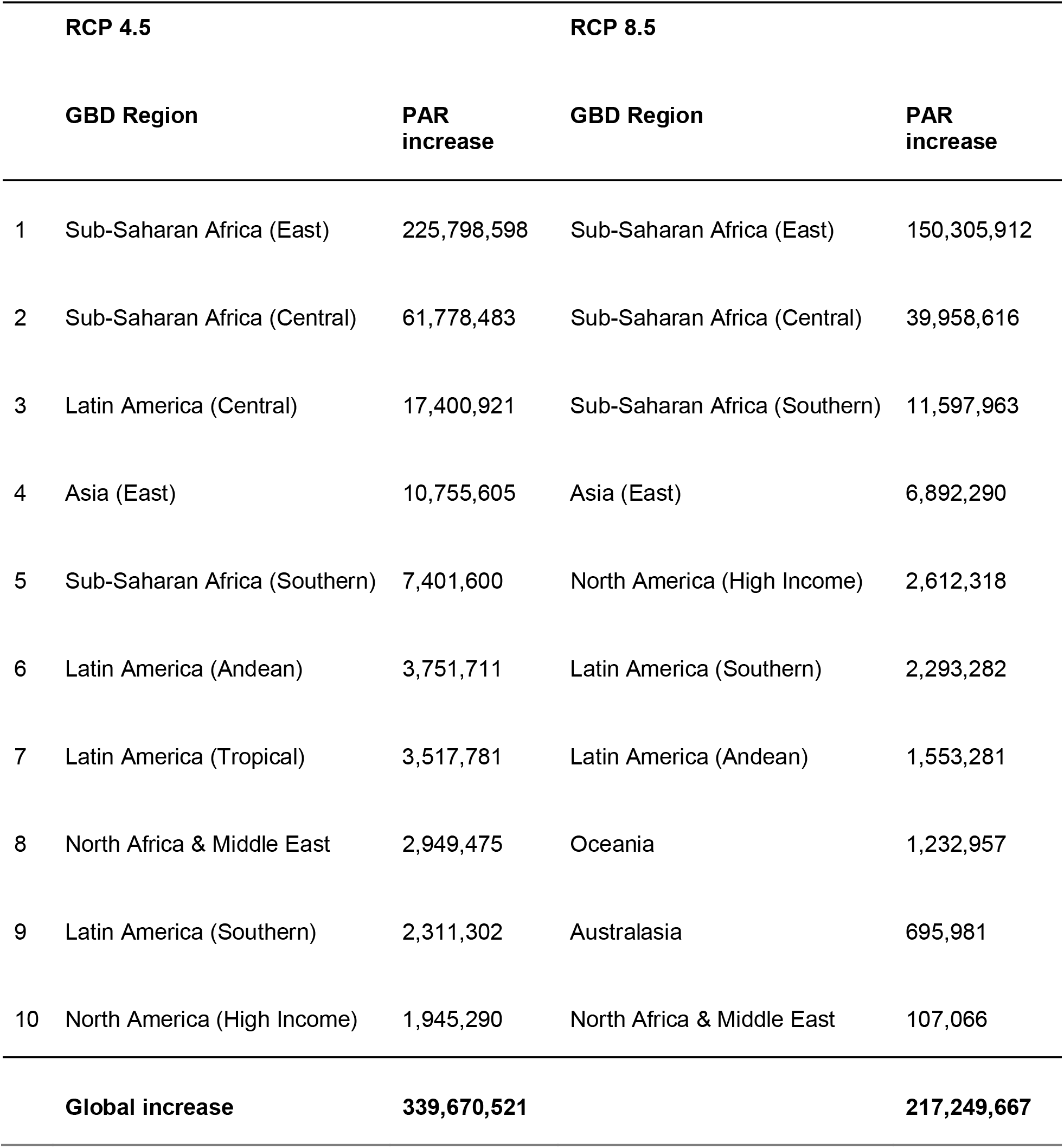
Top 10 Global Burden of Disease defined regional increase in people at risk (PAR) for year-round transmission suitability of *P. vivax* by *An. stephensi* in 2050, under RCP 4.5 (SSP2 population projection) and RCP 8.5 (SSP5 population projection) future climate scenarios, averaged across four general circulation models (GCMs) as described in the main methods. Global increase is the sum of all gains in PAR increases across all GBD regions.

**Table S3.** Top 10 Global Burden of Disease defined regional increase in people at risk (PAR) for one or months transmission suitability of *P. falciparum* by *An. stephensi* in 2050, under RCP 4.5 (SSP2 population projection) and RCP 8.5 (SSP5 population projection) future climate scenarios, averaged across four general circulation models (GCMs) as described in the main methods. Global increase is the sum of all gains in PAR increases across all GBD regions.

| RCP 4.5 |  | RCP 8.5 |  |
| --- | --- | --- | --- |
| GBD Region | PAR increase | GBD Region | PAR increase |
| 1 Asia (South) | 465,096,731 | Asia (South) | 187,123,365 |
| 2 Sub-Saharan Africa (East) | 283,814,459 | Sub-Saharan Africa (East) | 171,277,189 |
| 3 Sub-Saharan Africa (West) | 271,935,016 | Sub-Saharan Africa (West) | 165,798,742 |
| 4 North Africa & Middle East | 154,006,142 | Europe (Western) | 152,051,451 |
| 5 Europe (Western) | 75,893,897 | North America (High Income) | 147,947,883 |
| 6 North America (High Income) | 67,415,234 | North Africa & Middle East | 87,744,795 |
| 7 Asia (Southeast) | 50,393,023 | Sub-Saharan Africa (Central) | 25,042,405 |
| 8 Sub-Saharan Africa (Central) | 48,162,272 | Australasia | 23,093,746 |
| 9 Latin America (Central) | 44,930,762 | Latin America (Central) | 3,121,109 |
| 10 Australasia | 14,947,865 | Sub-Saharan Africa (Southern) | 2,914,570 |
| **Global increase** | **1,516,731,989** |  | **968,115,736** |

**Table S4.** Top 10 Global Burden of Disease defined regional increase in people at risk (PAR) for one or months transmission suitability of *P. vivax* by *An. stephensi* in 2050, under RCP 4.5 (SSP2 population projection) and RCP 8.5 (SSP5 population projection) future climate scenarios, averaged across four general circulation models (GCMs) as described in the main methods. Global increase is the sum of all gains in PAR increases across all GBD regions.

| RCP 4.5 |  | RCP 8.5 |  |
| --- | --- | --- | --- |
| GBD Region | PAR increase | GBD Region | PAR increase |
| 1 Asia (South) | 465,270,699 | Asia (South) | 187,452,995 |
| 2 Sub-Saharan Africa (East) | 287,838,551 | Europe (Western) | 180,670,956 |
| 3 Sub-Saharan Africa (West) | 271,933,624 | Sub-Saharan Africa (East) | 175,513,403 |
| 4 North Africa & Middle East | 154,326,951 | Sub-Saharan Africa (West) | 165,797,409 |
| 5 Europe (Western) | 104,894,102 | North America (High Income) | 150,498,663 |
| 6 North America (High Income) | 69,688,688 | North Africa & Middle East | 88,097,243 |
| 7 Asia (Southeast) | 50,497,397 | Sub-Saharan Africa (Central) | 25,400,517 |
| 8 Sub-Saharan Africa (Central) | 48,520,385 | Australasia | 23,311,798 |
| 9 Latin America (Central) | 43,071,846 | Latin America (Central) | 6,272,702 |
| 10 Australasia | 15,142,386 | Sub-Saharan Africa (Southern) | 2,926,014 |
| **Global increase** | **1,554,262,423** |  | **1,011,067,047** |

